# Predicting Gene Expression from DNA Sequence using Residual Neural Network

**DOI:** 10.1101/2020.06.21.163956

**Authors:** Yilun Zhang, Xin Zhou, Xiaodong Cai

## Abstract

It is known that *cis*-acting DNA motifs play an important role in regulating gene expression. The genome in a cell thus contains the information that not only encodes for the synthesis of proteins but also is necessary for regulating expression of genes. Therefore, the mRNA level of a gene may be predictable from the DNA sequence. Indeed, three deep neural network models were developed recently to predict the mRNA level of a gene directly or indirectly from the DNA sequence around the transcription start side of the gene. In this work, we develop a deep residual network model, named ExpResNet, to predict gene expression directly from DNA sequence. Applying ExpResNet to the GTEx data, we demonstrate that ExpResNet outperforms the three existing models across four tissues tested. Our model may be useful in the investigation of gene regulation.

## Introduction

For a cell in a living organism to function properly, necessary proteins must be synthesized at the proper time. All cells regulate the synthesis of proteins from information encoded in their DNA through transcription of genes into mRNAs and their subsequent translation into proteins. Gene expression is thus controlled first and foremost at the level of transcription. Much of this control is achieved through the interplay between proteins that bind to specific DNA sequences and their DNA-binding sites or DNA motifs. Transcription factors can bind to their DNA motifs in the promoter region of a gene to enhance or suppress the transcription of the gene. Epigenetic mechanisms such as DNA methylation can also regulate gene expression [1]. Studies have shown that DNA-binding factors and *cis*-acting DNA motifs play an important role in shapping DNA methylation [2, 3], which may in turn affect the expression of certain genes. The steady-state mRNA level of a gene is also affected by the degradation rate of the mRNA. The regulation of mRNA degradation is mediated by mRNA-binding proteins and non-coding RNAs such as microRNAs [4]. These mRNA-binding proteins and microRNAs bind to mRNA at specific binding sites. These mechanisms of regulating gene expression all allude that the information in DNA sequence upstream and downstream of the transcription start site (TSS) of a gene may determine the expression level of the gene. Such information may be exploited to build a computational model to predict gene expression levels.

Recently, three deep neural network models have been developed to predicted gene expression levels from DNA sequences [5, 6, 7]. The ExPecto framework employs a convolutional neural network, consisting of 7 convolution layers, 2 linear layers, and other layers such as pooling and ReLU layers, to predict 2,002 epigenomic features, including genome-wide histone marks, TF-binding and chromatin accessibility profiles, from a DNA sequence of 40 kbps centered at the TSS of a gene. Then, these 2,002 features are used by a linear model to predict the mRAN level of the gene. The Basenji architecture [6] consists of 12 convolution layers and other layers such as pooling and ReLU layers, and it predicts mRNA level of a gene directly from the DNA sequence of 131 kbps long. The Xpresso model [7] is composed of two convolution layers, two fully connected layers, and other layers such as pooling layers, and it predicts the expression level of gene from a DNA sequence of 10.5 kbps (7 kbps upstream and 3.5 kbps downstream of the TSS) and several mRNA features.

It is known that deep neural networks are difficult to train and adding more layers may not necessarily improve performance. To overcome these problems, deep residual network (ResNet) was introduced [8]. In image recognition, ResNet can significantly improve prediction accuracy comparing traditional convolutional neural networks (CNNs) [8]. In this work, we develop a method of predicting gene expression using residual network (ExpResNet) directly from DNA sequence. Our results show that ExpResNet outperforms ExPecto, Basenji, and Xpresso.

## Methods

### 2.1 Data processing

The gene expression and single nucleotide polymorphism (SNP) data from the Genotype-Tissue Expression (GTEx) Project [9] and the genome sequence of HG19(GRCh37) [10] were used to train the residual network to predict gene expression. The RNA-seq data containing the expression values of 20, 805 genes in 218 tissues of 450 individuals were downloaded from the GTEx Portal using the GTEx Analysis V6p release (dbGaP accession phs000424.v6.p1). For each gene, we used the nucleotide sequence, starting from 70,000 base pairs (bps) upstream of the transcription start site (TSS) to 25,000 bps downstream of the TSS, to predict the expression level of the gene. We extracted the 95,000 pb long sequence of each gene from the GRCh37 genome. We used the 4-channel one-hot encoding scheme to encode the DNA sequence. Each sequence were represented by a 4 × *L* matrix, where *L* is the length of the sequence. Each channel encoded one of the four bases adenine (A), thymine (T), cytosine (C), and guanine (D). There are other letters in the DNA sequence; for example, “R” and “Y” represent a purine and a pyrimidine, respectively. The encoding scheme for all possible letters is shown in Table 1. We used the average expression value of each gene in a tissue, as also used in [5], to train a tissue-specific ResNet for expression prediction. More specifically, let *x*_*ij*_ be the RPKM value of the *i*th gene in a tissue, then the average expression value for the *i*th gene is 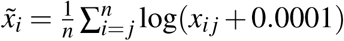, where *n* is the number of individuals.

**Table 1.** Encoding scheme for nucleotides

| A | T | C | G | - | . | N | R | Y | S | W | K | M | B | D | H | V |
| --- | --- | --- | --- | --- | --- | --- | --- | --- | --- | --- | --- | --- | --- | --- | --- | --- |
| 1 | 0 | 0 | 0 | 0 | 0 | 1/4 | 1/2 | 0 | 0 | 1/2 | 0 | 1/2 | 0 | 1/3 | 1/3 | 1/3 |
| 0 | 1 | 0 | 0 | 0 | 0 | 1/4 | 0 | 1/2 | 0 | 1/2 | 1/2 | 0 | 1/3 | 1/3 | 1/3 | 0 |
| 0 | 0 | 1 | 0 | 0 | 0 | 1/4 | 0 | 1/2 | 1/2 | 0 | 0 | 1/2 | 1/3 | 0 | 1/3 | 1/3 |
| 0 | 0 | 0 | 1 | 0 | 0 | 1/4 | 1/2 | 0 | 1/2 | 0 | 1/2 | 0 | 1/3 | 1/3 | 0 | 1/3 |

### 2.2 ExpResNet structure

ExpResNet is a deep residual network [8], and its structure is depicted in Figure 1. The model consists of five residual units, each followed by an adaptive average pooling layer, and two fully connected layers with a batch normalization a layer and a ReLU layer in between the two layers. Each residual unit have 4 input parameters: in is the number of input channels, out is the number of output channels, *k* and *d* are the kernel size and the dilation rate, respectively, used by the convolution layer inside the residual unit. Of note, the model structure described in Figure 1 was obtained after searching over a set of hyperparameters including the number of residual units, the number of layers in each residual unit, the number of kernels at each convolution layer, etc, as will be described later in the Training section. For illustration purpose, we present the final network structure here in the Methods section.

**Figure 1.**
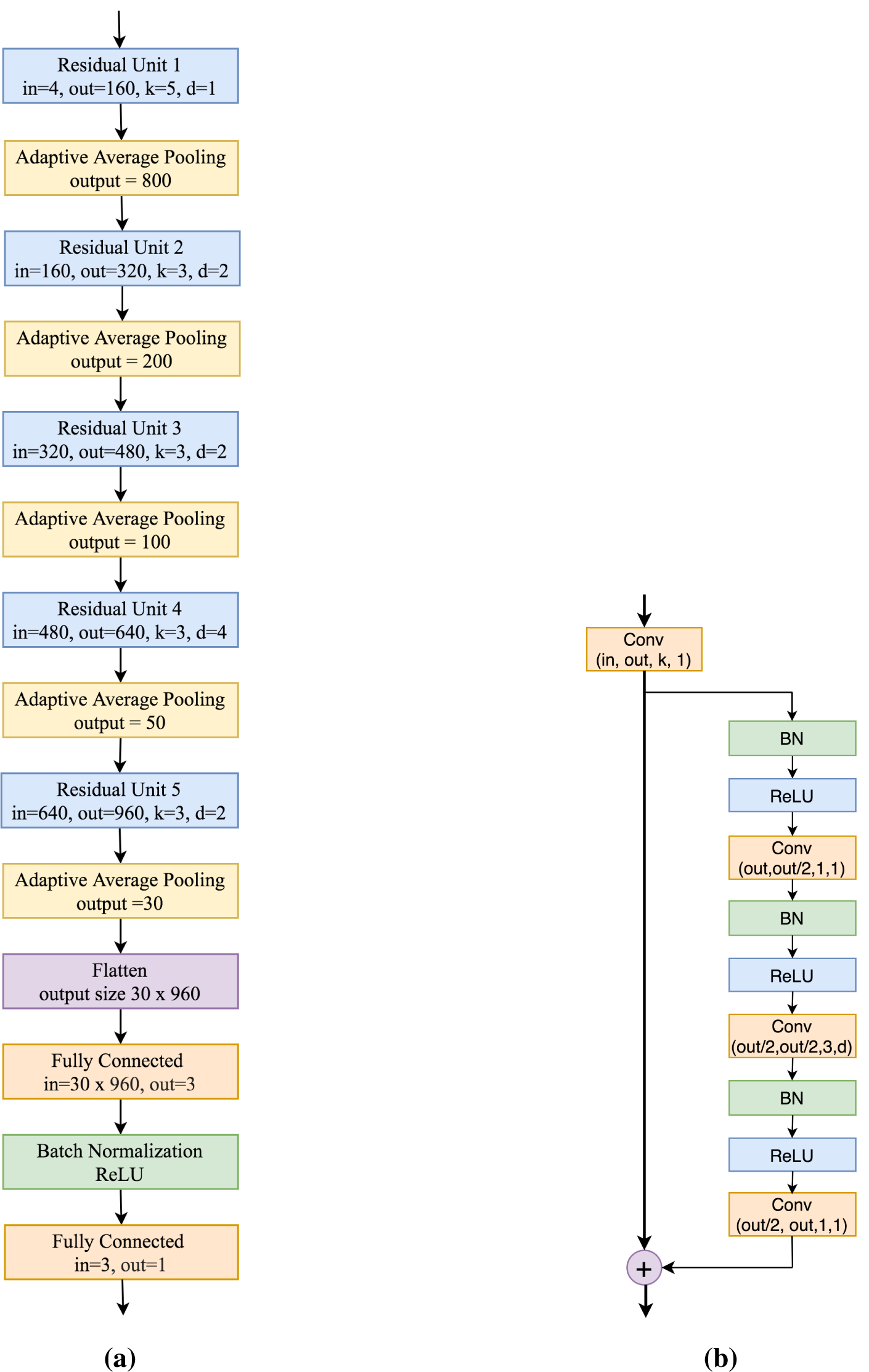
Block diagrams for the residual network (a) and the residual unit (b).

The structure of the residual unit is shown in Figure 1b. To increase the parameter efficiency, we adopted the design of the bottleneck residual unit [8], and used small convolutional kernel sizes suggested by the inception module [11]. The residual unit consists of an convolution layer followed by three submodules, each of which is composed of a batch normalization layer [12], an ReLU activation layer [13], and a convolution layer. The residual unit in [14] does not have the first convolution or weight layer; for ease of presentation, we here include the first convolution layer in the residual unit, although we can leave it outside of the residual unit. The stride of all convolution layers was set to 1. The first convolution raises the number of feature maps so that the number of the output channels of the residual unit is eventually larger than the number of input channels. At the convolution layer of the first submodule, the number of output channel is half of the number of input channels, and the kernel size is equal to 1. Therefore, this convolution layer essentially performs dimension reduction. At the convolution layer of the second submodule, the kernel size is 3. The number of input channels is equal to the number of the output channels for the second submodule. The last submodule uses 1 × 1 convolution with the number of output channel being twice the number of the input channel, which raises the dimensions back to be the same as the one at the input of the first submodule. Eventually, the residual unit does not change the number of channels and lengths of feature maps. Finally, the output of the final submodule is added to the input feature map of the residual unit as the output of the residual unit.

As described earlier, each of the five residual units starts with a convolution layer. The numbers of kernels of the first convolution layer in five residual units are 160, 320, 480, 640, and 960, with the kernel size equal to 5, 3, 3, 3 and 3, respectively. Following each residual unit, an adaptive average pooling operation is applied to down-sample the feature map. The parameter *output* for each pooling layer in Figure 1a specifies the dimension of the output features. The pooling window size was changed adaptively in different residual blocks so that the output dimension is equal to the value of parameter *output*. If the stride is smaller than the window size, there will be overlap between neighboring windows. For a given window size, we tested different strides, and found that when the stride was set equal to the window size to avoid the overlap between neighboring windows, better performance was achieved than the case where there was overlap between neighboring windows. Therefore, we chose the stride equal to the window size. The feature maps at the output of the last residual unit were flattened and fed into a fully connected layer that had 3 output nodes, connected to a batch normalization layer followed by a ReLU layer. Finally, the output of the final fully connected layer is a scalar, which gives the predicted expression value.

Of note, to increase the receptive field of the neural network to learn nonlocal representation of features in relatively long DNA sequence, we employed the dilated neural network structure [15].

In our residual network, we first increased the dilation rate and then decreased it to adapt to the “degridding” problem [16]. The dilation rates are hyperparameters, and the set of optimal values was found to be 1, 2, 4, 2, and 2, respectively, for five residual units.

### 2.3 Training

We split the whole dataset into a training set, a validation set and a test set. Genes on X or Y chromosomes were excluded from training and testing. The 1,017 genes on chromosome 8 were set as the test set. The 966 genes on chromosome 10 were used as the validation set, and the genes the remaining chromosomes serves as the training set. The mini-batch stochastic gradient descent (SGD) method was used for training the model. The batch size was chosen to be 8, and the initial learning rate was 0.00005, and an efficient SGD method Adam [17] was employed.The loss function is the mean square error at the output of the model. Early stopping [18] was used to avoid over-fitting the neural network. After every 100 iterations of the training, Pearson’s correlation coefficient (PCC) between the predicted and true gene expression values was evaluated on the validation dataset. The training would be terminated, if the PCC was lower than the minimum PCC in previous 3 consecutive iterations. We also used data augmentation to increase the data instances by shifting the DNA sequence by a size within 0-30 bps during each epoch of the training. The PyTorch platform [19] was used to train the neural network.

There are a number of hyperparameters that need to be chosen properly so the the model can offer good prediction power. The ExpResNet model described in Figure 1 was obtained after a set of hyperparameter values were tried. The core hyperparameters include: the number of layers, the kernel size and the number of kernels at each convolution layer, down-sampling configuration at each layer, the dilation rate, and the dimension of each convolution layer. The number of hyperparameters increases exponentially with the number of layers, which may incur a huge computational burden in searching for the set of optimal values for the hyperparameters.

The first convolution layer of the residual network will learn basic spatial features in the DNA sequence, while the later layers will learn more abstract spatial features. Therefore, the kernel size of the first convolution layer was chosen to be 5, and the kernel size of later convolutinal layers was smaller, equal to 3. We roughly divided the hyperparameters into two subsets: one contains the number of layers and the number of kernels that determine the learning capacity of the model, and the other mainly contains the down-sampling rate that controls the resolution of the model [8, 20]. The higher the level of abstraction of the features, the lower the resolution. During the hyperparameter searching process, we fixed the relationship among the number of kernels at the different layers and the relationship among the down-sampling rates at different pooling layers. Specifically, the number of kernels per layer was increased by a factor between 1.5-2. For example, in the ExpResNet model in Figure 1, the number of filters at five residual units are 160, 320, 480, 640, and 960, respectively. In this way, we could control the learning capacity of the neural network by only setting the number of kernels in the first layer and the total number of layers. The down-sampling rate per layer was increased by a factor close to 2 [8], and thus, the resolution at the first pooling layer will control the resolutions at all subsequent layers. We gradually increased the number of kernels at the first layer and the number of layers until the PCC on the validation dataset reached a plateau. For the hyperparameters in the second subset, they were tuned using random search [21] by trying a set values of the down-sample rates at the first pooling layer.

## Results

We used the GTEx data to compare the performance of Expecto [5], Expresso [7], Basenji [6], and our ExpResNet. The Expecto neural network trained with genomic sequences and epigenomic data from ENCODE [22] and Roadmap Epigenomics [23] projects was available. Following the steps in [5], we used the pretrained Expecto neural network to predict 400,400 epigenomic features from the DNA sequence of 40,000 bps, 20,000 bps upstream and 20,000 bps downstream of the TSS of a gene. The 400,400 epigenomic features were input to a spatial transform and reduced to 20,02 features. These spatially transformed features were then used to predict gene expression levels for each tissue with an *L*_2_-regularized linear regression model fitted by a gradient-boosting algorithm XGBoost [24]. Since we used the newest version of XGBoost which was different from the one used by Expecto in [5], the value for the hyperparameter of *L*_2_-regularization suggested by Expecto might not work well. Therefore, we used cross validation to search over a set of 81 values {10^−4+0.1*k*^, *k* = 0, 1,, 80}, and found the optimal value for the hyperparameter, which was then used to estimate the linear model for predicting gene expression levels from the 20,02 spatial features. Another widely used regularized regression is the elastic net [25], which uses the regularization term 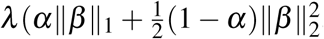, where *λ* > 0 and 0 ≤ *α* ≤ 1 are two hyperparameters, *β* is a vector containing regression coefficients, and ∥ · ∥_*q*_, *q* = 1, 2, stands for *q*-norm. Software package glmnet [26] was used to estimate elastic net models. Ten-fold cross-validation was employed to search over a set of 30 values that were automatically generated by glmnet for the optimal value of *λ* and over the set {0, 0.01, 0.05, 0.1, 0.3, 0.5, 0.7, 0.9, 0.95, 0.99, 1} for the optimal value of *α*. The final model parameters in *β* were obtained from the training data with the optimal values of *λ* and *α*, and then were used to predict gene expression values in the test dataset. We reported the results obtained with XGBoost and glmnet as Expecto(xg) and Expecto(gl), respectively. For Expresso, we followed the specifications of the final model architecture [7], implemented the model using PyTorch [19], and predicted the expression value of a gene with the model from the DNA sequence of 10,5000 bps, 7,000 bps upstream and 3,500 bps downstream of the TSS of the gene. For Basenji, we used model architecture and the hyperparameters specified at the authors’ website (https://github.com/calico/basenji), and replaced the loss function with the mean square error loss for prediction of gene expression, and connected the second last layer to the output with a linear layer. The Basenji model predicts gene expression from a DNA sequence of 131,000 bps, 65,500 bps upstream and 65,5000 bps downstream of the TSS of the gene. We predicted gene expression values in four tissues: breast mammary tissue, muscle skeletal tissue, ovary tissue, and whole blood. After expression values of genes in the test dataset were predicted, Pearson’s correlation and Spearman’s correlation between the predicted values and true values were calculated as the performance measure.

The performance of the four neural network models in prediction of gene expressions in four tissues is given in Table 2. It is seen that our ExpResNet offers the best performance across all four tissues. For fair comparison, we also applied ExpResNet to the same DNA sequences that were used by Expecto and Xpresso. Our ExpResNet again outperforms Expecto and Xpresso. These results clearly show that the residual network improves the prediction accuracy over other convolutional neural networks.

**Table 2.** Performance of four neural network models in prediction of gene expression

| Methods | Sequence length (kbps) |  | Pearson's/Spearman's correlation |  |  |  |
| --- | --- | --- | --- | --- | --- | --- |
|  | upstream | downstream | Breast | Muscle | Ovary | Whole blood |
| ExpResNet | 70 | 25 | 0.7954 / 0.7911 | 0.7913 / 0.7939 | 0.8026 / 0.8016 | 0.7502 / 0.7536 |
| Basenji | 65.5 | 65.5 | 0.5230 / 0.5665 | 0.5369 / 0.5776 | 0.5256 / 0.5646 | 0.5824 / 0.6108 |
| Xpresso | 7 | 3.5 | 0.7443 / 0.7236 | 0.7400 / 0.7221 | 0.7687 / 0.7450 | 0.7073 / 0.7085 |
| ExpResNet | 7 | 3.5 | 0.7642 / 0.7631 | 0.7574 / 0.7512 | 0.7845 / 0.7742 | 0.7456 / 0.7460 |
| Expecto (xg) | 20 | 20 | 0.7336 / 0.7401 | 0.7340 / 0.7363 | 0.7488 / 0.7502 | 0.7097 / 0.7187 |
| Expecto (gl) | 20 | 20 | 0.7378 / 0.7457 | 0.7388 / 0.7416 | 0.7496 / 0.7524 | 0.7089 / 0.7184 |
| ExpResNet | 20 | 20 | 0.7725 / 0.7706 | 0.7917 / 0.7844 | 0.7795 / 0.7870 | 0.7526 / 0.7623 |

## Conclusion

In this work, we developed a residual network model for predicting gene expression form DNA sequence. Using the genomic and gene expression data of four tissues in the GTEx database, we demonstrated that our deep neural network model outperformed three existing convolutional neural network models.

## Notes

### Competing Interest Statement

The authors have declared no competing interest.

